# Image classification and cognition using contour curvature statistics

**DOI:** 10.1101/2021.08.25.457634

**Authors:** Andrew Marantan, Irina Tolkova, L. Mahadevan

## Abstract

Although the higher order mechanisms behind object representation and classification in the visual system are still not well understood, there are hints that simple shape primitives such as “curviness” might activate neural activation and guide this process. Drawing on elementary invariance principles, we propose that a statistical geometric object, the probability distribution of the normalized contour curvatures (NCC) in the intensity field of a planar image, has the potential to represent and classify categories of objects. We show that NCC is sufficient for discriminating between cognitive categories such as animacy, size and type, and demonstrate the robustness of this metric to variation in illumination and viewpoint, consistent with neurobiological constraints and psychological experiments. A generative model for producing artificial images with the observed NCC distributions highlights the key features that our metric captures and just as importantly, those that it does not. More broadly, our study points to the need for statistical geometric approaches to cognition that build in both the statistics and the natural invariances of the sensory world.

---

Humans and other primates are adept at recognizing objects within a visual field remarkably quickly and accurately, and forming neural representations that are spatially distributed in the inferotemporal cortex into general cognitive categories such as animacy, size, type (faces vs places) etc. How this happens is still not well understood. There are at least two approaches to addressing this question: via a bottom-up approach to understanding the neural areas and circuits responsible by systematic microscopic studies, or by taking a top-down approach that ignores the details takes a perspective that is deliberately a high-level approach to solving the inverse problem: what kind of image representation could agree with neural activation patterns measured in participants in response to different aspects of chosen image features and stimuli [1]? Eventually, we hope to be able to merge these levels of understanding into a unified whole that accounts for the natural statistics of categories in the visual world, and how the evolution, and development of the visual cortex as it learns to compress, classify and comprehend the external environment.

At a high level, visual processing in primates is roughly divided across the ventral and dorsal pathways, with the former responsible for characterizing objects in the visual field (“what”), and the latter for guiding interactions with the objects (“how”) [2, 3]. Within the ventral stream, object recognition is known to occur in the inferotemporal (IT) cortex [4, 5]. Studies of fMRI in both humans and non-human primates in response to viewing different stimuli have uncovered multiple cognitive categories or dimensions — such as size (from small to large objects), animacy (from animals to inanimate objects), body parts (such as faces, hands, bodies) — which elicit neural activation in different spatial domains of the IT cortex [6–9]. At the neural level, studies have iteratively simplified stimuli to recover “critical features” which maximally activate a cell [10, 11]. In some regions of the visual pathway, such features permit intuitive explanations – for instance, the middle temporal (MT) cortex and middle superior temporal (MST) cortex are known to represent visual motion through a collection of neurons encoding direction and speed [12, 13]. Interestingly, within the intermediate subregions of the ventral cortical pathway leading to the IT cortex – areas V1, V2, and V4 – the critical features associated with images of varying contours are found to represent both simple properties, such as position and orientation, but also a higher-order property: curvature [14–17], and metrics based on this quantity are strongly correlated with neural dynamics [18, 19].

In psychology and psychophysics, the significance of contour curvature in perception has been suggested at least since the 1950s [20], when it was suggested that perceptual information along a shape boundary is not equally distributed, but rather concentrated in regions of high curvature — a proposal has since been supported both empirically and through an information-theoretic framework [21, 22]. In particular, studies have shown that we exhibit hyper-acuity in the perception of curved lines [23–25]; in fact studies have found that the mechanisms for discriminating contour curvature are selective for spatial frequency and orientation, agreeing very closely with studies of neural features [26]. Furthermore, given the evidence for orientational selectivity in the visual cortex, curvature which is the spatial rate of change of orientation is a relatively easy feature to extract. In machine vision, while the notion of curvature has been used widely used [27–32], most image descriptor for object recognition and classification tasks are composed of histograms of pixel-based metrics – most notably, the histogram of oriented gradients (HoG) approach [33]. While several studies have combined HoG with locally binned histograms of curvature to show improved performance in numerical tasks, they use curvature only to augment existing methods [34, 35], rather than as a distinct metric. A promising alternative is that of curvature scale space (CSS) - a representation of shape through contour curvature calculated at different magnitudes of Gaussian smoothing [36] that has proved efficient and successful in problems of corner detection, clustering, shape indexing and retrieval, and silhouette-based object recognition [37–40]. However, while CSS is rooted in and motivated by the mathematical invariances desirable for shape analysis, it does not seem to have been extended to quantifying two-dimensional images, or to our understanding of biological perception. Finally and most recently, while neural networks have been successful at solving many problems in computer vision [41], and are promising models of biological processing [42, 43], they do not usually provide an interpretable understanding of the intermediary image representations that underlie perception.

Given these insights from neurobiology, psychology, and computer vision, how might one construct a computationally meaningful and interpretable image descriptor? Here, guided by the need for an invariant description to Euclidean motions, we propose the use of a simple statistical geometric measure, the “normalized contour curvature” (NCC) distributions: a probability distribution of curvatures within smoothed natural images. We construct NCC through pooling of nonlinear transformations of an image’s contour curvature content – a simple calculation which has plausible implementability within neural circuitry, emphasizing that it is important to think of a statistical measure of this geometric quantity given the nature of noise in images. We show that NCC satisfies desired properties of shape, interpret calculations over example stimuli, and demonstrate that this metric carries sufficient informational content for distinguishing between cognitive categories. Finally, we derive a generative model for constructing images corresponding to a given NCC distribution to help us understand when it works and perhaps more importantly, when it does not work.

## CONTOUR CURVATURE COMPUTATION

For the compression, classification and comprehension of images treated as shapes, any meaningful perception should satisfy some basic invariances that include (i) Global Translation Invariance:, i.e. shape is independent of location in space (ii) Global Rotation Invariance:, i.e. shape is independent of orientation (iii) Resizing/ Scale Invariance:, i.e. shape is independent of overall scale (iv) Image Representation Invariance:, i.e. shape is independent of rescaling the intensity map. While there are known cognitive exceptions (e.g. squares/diamonds, upside-down faces etc.), these are specialized and we will not consider them here.

For an image that is characterized by a two-dimensional intensity field, constant intensity contours typically form closed contours. In a smooth differential-geometric setting, the curvature of these curves is invariant to global translation and rotation and thus forms a natural candidate for an invariant description. Letting *f* (*x, y*) represent the intensity at pixel (*x, y*) and *f_x_, f_y_, f_xx_, f_yy_* and *f_xy_* represent the first and second derivatives, we can write the contour curvature (CC) at point (*x, y*) as

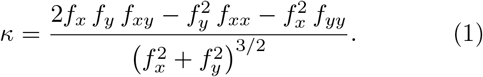

We note that this approach is different from using the intensity values of a given image to describe a height map and then compute the curvature tensor of the surface and thence the Gaussian or mean curvature at each pixel [44], although the two are of course related. Furthermore, we note that calculation of curvature follows naturally from orientational information that the retina is well known to respond to; curvature is just the spatial variation in orientational information and can be deduced approximately via a differencing scheme that is analogous to a difference of Gaussians. Though the contour curvature is invariant under translation, rotation and intensity scaling, it is not invariant to scale changes. To overcome this issue, we take the largest dimension of an image to be of unit length, so that the curvature of a circle fitting just inside the (square) image has radius *r* = 1/2 and hence curvature *κ* = 2. Finally, it is numerically useful to map the contour curvature defined on the whole real line to a finite interval which we choose to be [−1,1]. Thus we define the normalized contour curvature (NCC) as

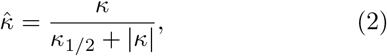

The parameter *κ*_1/2_ sets the value of *κ* which maps to 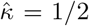, and makes the NCC easier to interpret; by taking *κ*_1/2_ = 2, a circle with curvature *κ* = 2 (i.e. a circle inscribed in a square image) is mapped to 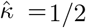 (Fig. 1). Note that this definition corresponds to a closed-form inverse relation.

**FIG. 1.**
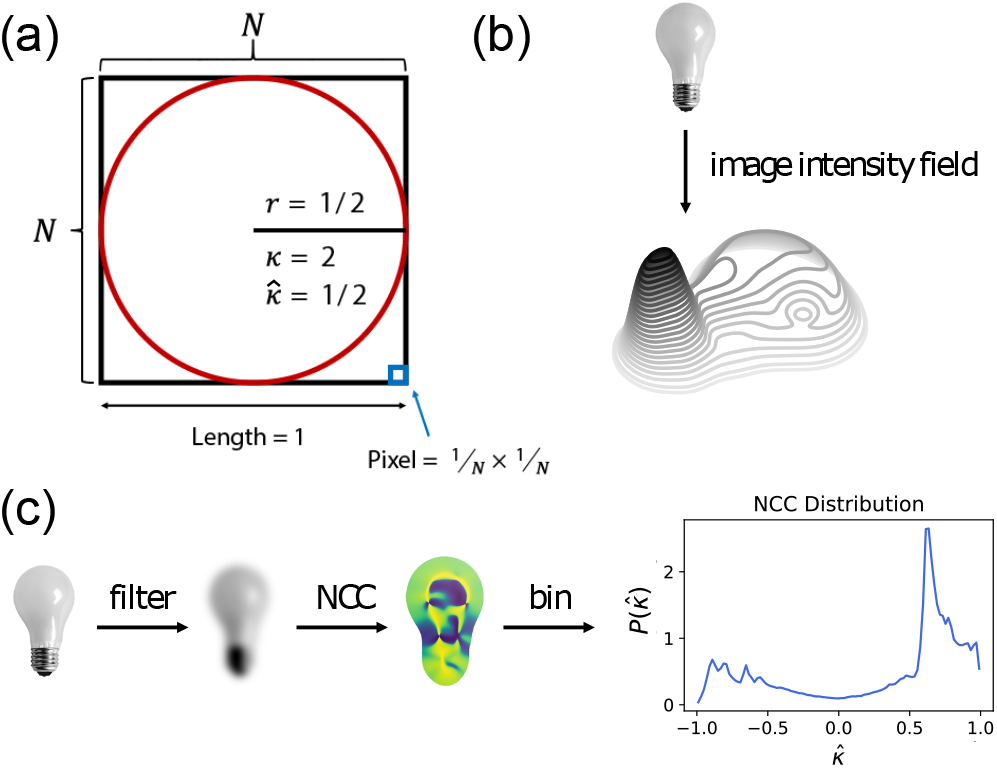
(a) The NCC is defined (by Eq. 2) such that the NCC along a circle inscribed in a square of side length 1 is mapped to 1/2. **(b)** When calculating NCC, we consider the level curves of a 3D surface defined by the pixel intensities of an image. This figure shows the level curves for the light-bulb image in (c). **(c)** This sequence shows the pipeline for calculating NCC. Starting with an image of a lightbulb, we apply Gaussian smoothing, calculate NCC for each pixel following Eq. 2, and finally histogram the values to result in a probability density.

In a discrete computational setting, pixel intensities can be used to construct level sets of constant intensity and compute the curvature of these contours at every pixel. While rescaling the intensity map does change the intensities of the contours, it does not change their overall shape, and so this method is manifestly invariant under intensity rescaling. To get a smooth surface interpolant from the image intensities, we filter the image using a Gaussian kernel that has zero mean and a standard deviation *ρ* and thus avoid the computational task of constructing the contours passing through each pixel and calculate the contour curvature directly in terms of numerical derivatives [45] of the filtered image intensity, and directly apply Eq. 1. This produces an “image” of the contour curvature, which can be converted to the normalized contour curvature via Eq. 2.

Finally, we use the normalized contour curvature image to construct a histogram for the original image, which we then convert to a probability density to produce the NCC distribution. We use equally-spaced bins spanning from 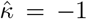 to 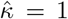, choosing an odd number of bins in order to have a bin centered on 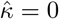. In order to count only curvatures corresponding to the object in the image, we ignore pixels corresponding to background elements. This image processing pipeline along with some examples is shown in Fig. 2 (see also Supplmentary Information - SI, Section A).

**FIG. 2.**
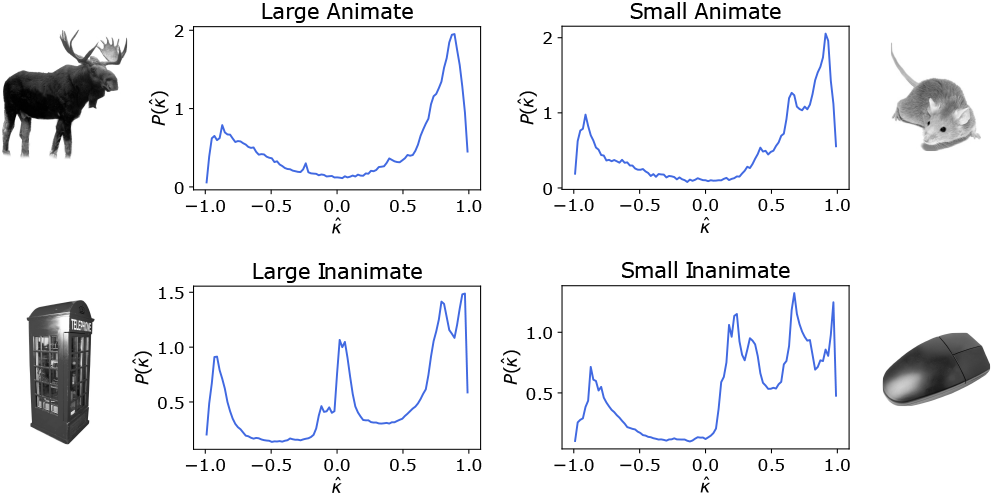
NCC probability densities calculated for example images from the stimulus dataset presented in [7], across four categories: a moose (large animate), mouse (small animate), telephone booth (large inanimate), computer mouse (small inanimate). Notice that the prevalence of straight lines in the telephone booth and computer mouse results in peaks near 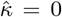, and that both mice contain higher probability mass in the low positive values (around 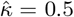). We find both of these features to be characteristic of animacy and size (Fig. 4).

## BAYESIAN IMAGE CLASSIFICATION OF ANIMACY AND SIZE

To evaluate the role of NCC as an image classifier, we investigate whether the NCC metric carries sufficient informational content for a simple binary classifier to distinguish between different cognitive categories. Our data were drawn from those used in fMRI studies on human participants who were shown real images distinguished by features such as animacy and size [7] that led to spatially localized responses in the IT cortex, and various artificial image sets [46] titled “texforms” – which, while designed to be unrecognizeable, preserved enough low-level structure to elicit a similar neural response to the images from which they were generated.

### Methods

For the natural images, the stimulus set contained 60 objects per each of four subcategories [7] – small animate, small inanimate, large animate, large inanimate – for a total of 240 images. The animate images spanned the phylogenetic tree, including mammals, reptiles, birds, and fish; the inanimate objects featured everyday items varying from a thimble to a firetruck. In all images, objects were centered on white background.

To classify a given image into one of two predefined classes *C*_1_ or *C*_2_, we use a slightly-modified log-likelihood scheme that relies on the supervised learning of two probability distributions, 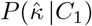 and 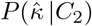, representing the probability for a pixel in a given image to have normalized curvature 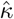 given that it belongs to either *C*_1_ or *C*_2_. In practice, we bin the normalized curvatures, and the aforementioned distributions become probability vectors: 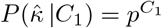 and 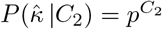, where the *n*th element 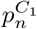 (or 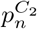) describes the probability for 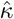 to be in the *n*th bin.

To construct *p*^*C*_1_^ (or *p*^*C*_2_^), we simply calculate NCC histograms for all images in the *C*_1_ (or *C*_2_) training set, add them together, and normalize by the total number of counts. Then, we can classify an image by calculating its NCC distribution *p_n_*, dividing by the total number of counts to obtain *q_n_*, and compute the log-likelihood:

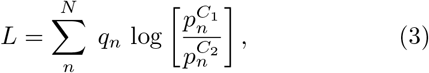

where *N* is the number of bins. If *L* > 0, the image is classified as belonging to *C*_1_; is *L* ≤ 0, it is classified as belonging to *C*_2_. We note that *L* can also be thought of as the difference between the Kullback-Leibler (KL) divergence of *q* from *p*^*C*_1_^ and the KL divergence of *q* from *p*^*C*_2_^. Although we only use the sign of *L* for predicting the category of an image, the magnitude of *L* can inform the classification likelihood; large values of |*L*| are linked with a higher confidence of classification, while values with |*L*| ~ 0 have a lower confidence.

In addition to analyzing natural images, we also considered the paired image/texform stimulus set introduced in [46], which contains a dataset of 120 objects (30 from each of the four size/animacy sub-categories), along with corresponding texforms. We perform the same preprocessing steps as described previously, with one difference in the processing of the background, whereby we take advantage of the provided green-screen variant to isolate the background pixels and set them to white to match prior analysis; applying this same “background mask” introduces an outline and background to the texform.

### Natural image classification

To test the efficacy of the normalized contour curvature as an image statistic and visualize the distinctions between classes, we plot the mean and variance of NCC distributions for each class in Figure 4. We highlight two important features. First, unlike animals, inanimate objects have a high prevalence of straight lines and edges, leading to a peak at 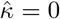. Second, small objects contain a higher density of intermediate curvature values, while the NCC of larger objects is more heavily concentrated at the ends of the distribution. This is likely because the characteristics and details of small objects are proportionately larger, resulting in lower absolute curvatures, while the fine-scale detail of images of large objects results in higher absolute curvatures.

**FIG. 3.**
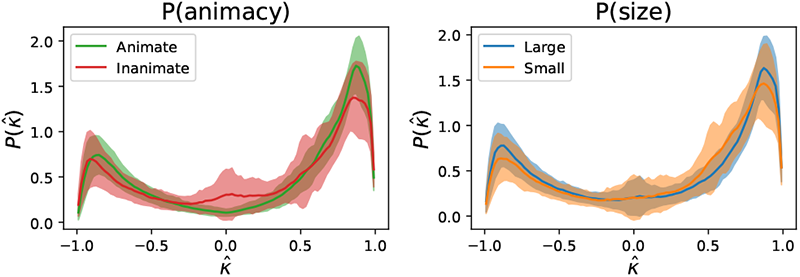
The solid line indicates mean NCC probability densities for each of the four image categories from the stimulus dataset presented in [7], and the shaded region indicates one standard deviation. We see that NCC for the inanimate category is characterized by high density at 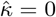 (representing the amount of straightness in the image) and NCC for the small category is characterized by slightly higher probability density in intermediate positive NCC values (around 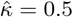).

**FIG. 4.**
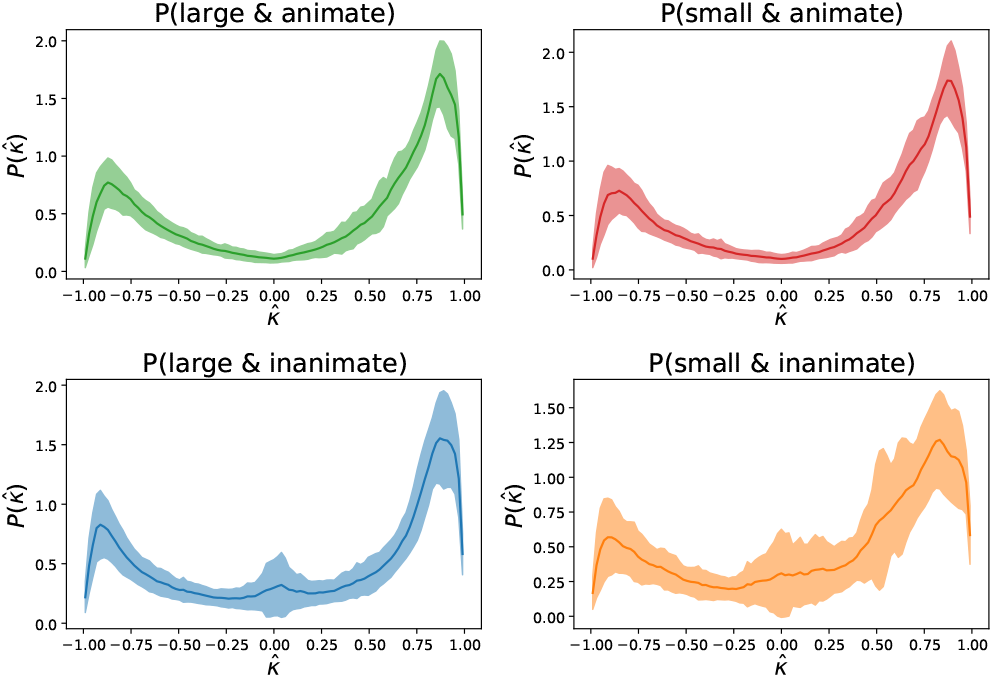
Here, we consider distributions for each of the four sub-categories of images from the stimulus dataset presented in [7], separated both by size and by animacy. We observe similar characteristics as described in Figure 3, though find that the distinction between large and small is much more prominent for inanimate images, which is consistent with the tripartite cognitive organization found in [7].

To classify animacy over both large and small objects, and classify size on both animate and inanimate objects, we ran 1000 randomized trials for both tasks, adhering to a 30%/70% training/testing split. Our aggregate results are shown in Figure 5. From the comparison of true positive and false positive rates in Figure 5, we see that these distinctions are sufficient for a simple Bayesian classifier to distinguish animacy within both large and small objects, and size within inanimate objects (with poorer performance on classifying size within animals). But in fact, this general tripartite organization – of small objects, animals, and large objects – is consistent with studies of neural activation within the occipito-temporal cortex for human observers [7].

**FIG. 5.**
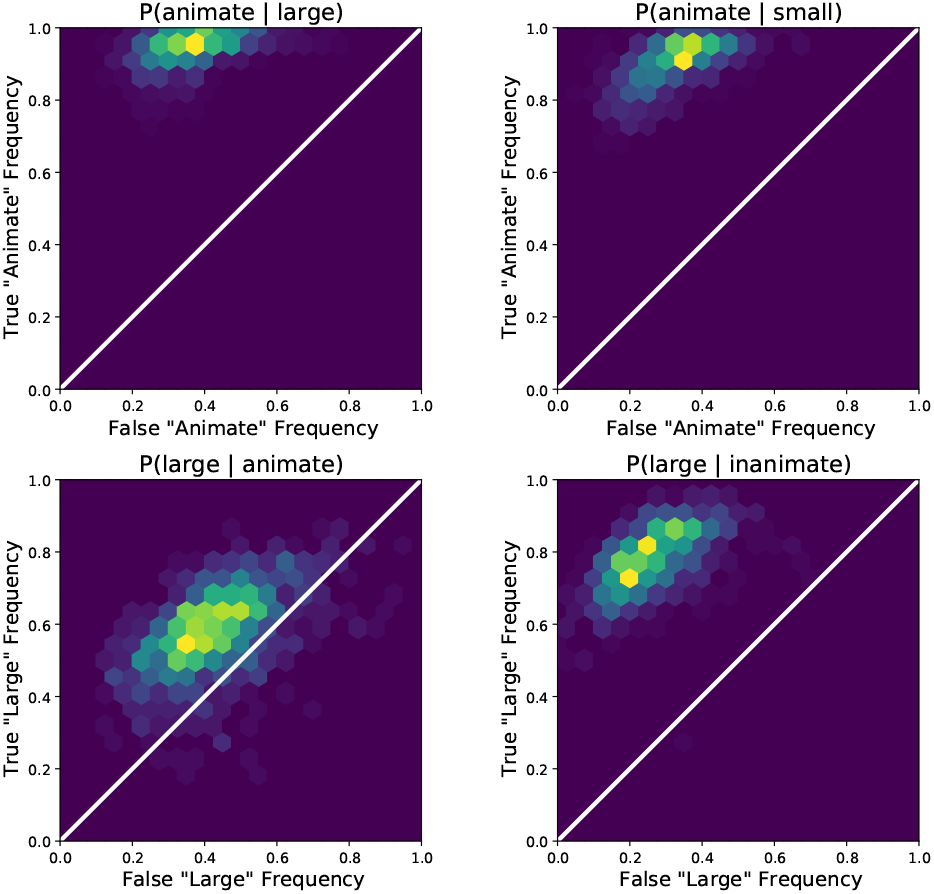
We examine the accuracy of a binary classifier (Eq. 3) by visualizing 2D histograms of the false positive rate against true positive rate over 1,000 randomized trials. Classification is performed separately across the four categories for the stimulus dataset presented in [7]; for instance, the top right histogram shows results for a binary classification tasks in which images with large objects are classified as animate or inanimate. We find that it is significantly more difficult to differentiate large animate and small animate images, while the other categories have high classification accuracy. These results are consistent with the tripartite domain separation observed in [7] across large inanimate, small inanimate, and animate objects.

The failure of the classifier exposed via misclassified samples, i.e. false positives and false negatives for each of the four classification tasks are shown in Fig. 6. Interpreting these mistakes in the NCC-based classifier is illuminating. For instance, in classifying large objects by animacy, most errors occur in inanimate objects with thin protruding “appendages”. This is likely because the thin components fade due to Gaussian blurring, resulting in high magnitude curvatures at their end-points, which align more closely with the animate distributions than with characteristically (straight) curvature-free inanimate distributions. When classifying small objects, most errors occur in inanimate objects which resemble animacy, such as a pinecone, and floral-patterned household items, all of which have rounded features. On the other hand, the snail is a false negative, suggesting that the shell is uncharacteristic of animate images. In this context, we find that the most difficult task is distinguishing size within animate images: misclassified samples include both large animals with the rounded shape usually associated with small objects, and small animals of more irregular and elongated shapes. Inanimate objects can more successfully be separated by size, and the mistakes often correspond to box-like large objects and elongated small objects.

**FIG. 6.**
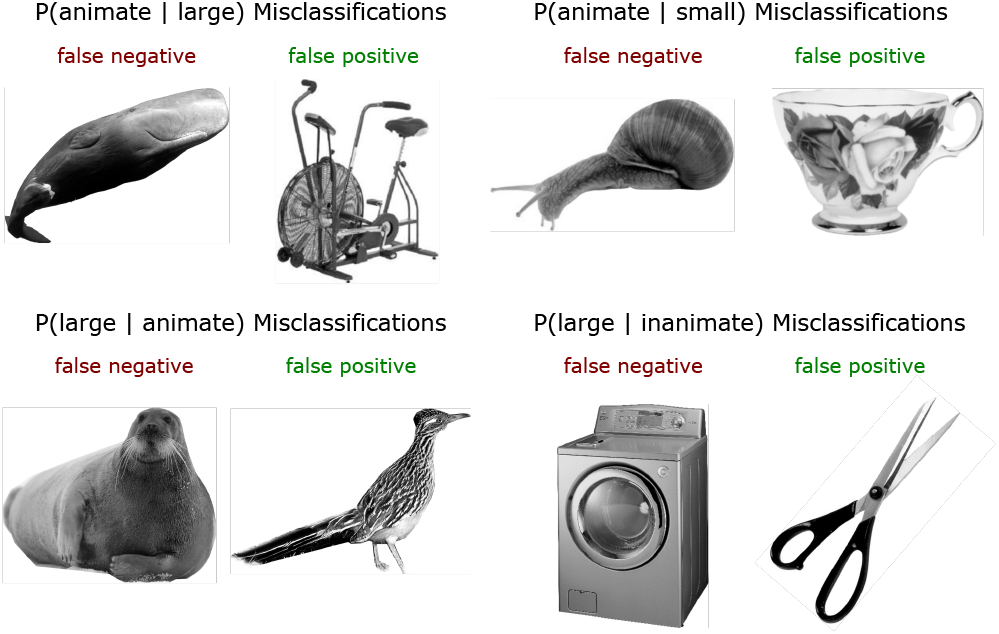
To understand the sources of error within our classifier (Eq. 3), we consider and interpret misclassified images. Specifically, we show a false negative and a false positive sample for each of the four classification tasks. We find that these mistakes are consistent with our observed differences in large/small and animate/inanimate NCC distributions. Since small objects are characterized by a peak in intermediate positive curvature, large objects are often misclassified when they have a rounded form, and small objects are often misclassified when comprising an elongated or irregular form. On the other hand, inanimate objects can be misclassified due to textured structure or when containing ‘‘appendages”. Animate objects are rarely misclassified as inanimate.

All our results so far use a particular value of the Gaussian blurring filter to remove high frequency information, namely *ρ* = 0.04 and a training fraction of 30%. Classification success rates exhibit some dependence on *ρ* and a weak dependence on the training fraction; performance suffers when *ρ* is very small (image noise dominates) and when *ρ* is very large (image details begin to be entirely washed out), but the success rates are not sensitive to small changes in *ρ* near the optimally-performing value (see SI-Section B and Figs. S2, S3, S4 for visualizations of these dependencies).

### Texform image classification

Images of natural objects are distinguished by their correlation statistics. Texforms attempt to scramble these [46] and thus serve as a test of our binary Bayesian classifier for distinguishing animacy and size. In Figure 7, we show that texforms preserve similar features as natural images, and moreover, have very similar classification result. The top row of shows the NCC distributions for texforms across both animacy and size. While texforms do not retain the large straight or flat regions characteristic of inanimate images, we do see slightly higher zero-curvature content in the inanimate distribution, and find that this difference is sufficient for classifying animate from inanimate texforms with relatively high confidence. In addition, the primary difference between large and small texforms is in intermediate positive curvatures, as in natural images. The bottom row of Figure 7 shows the true positive / false positive results from 1000 trials of a Bayesian classifier (see SI, Section B, and Figure S5 for results across all four animacy/size subcategories).

**FIG. 7.**
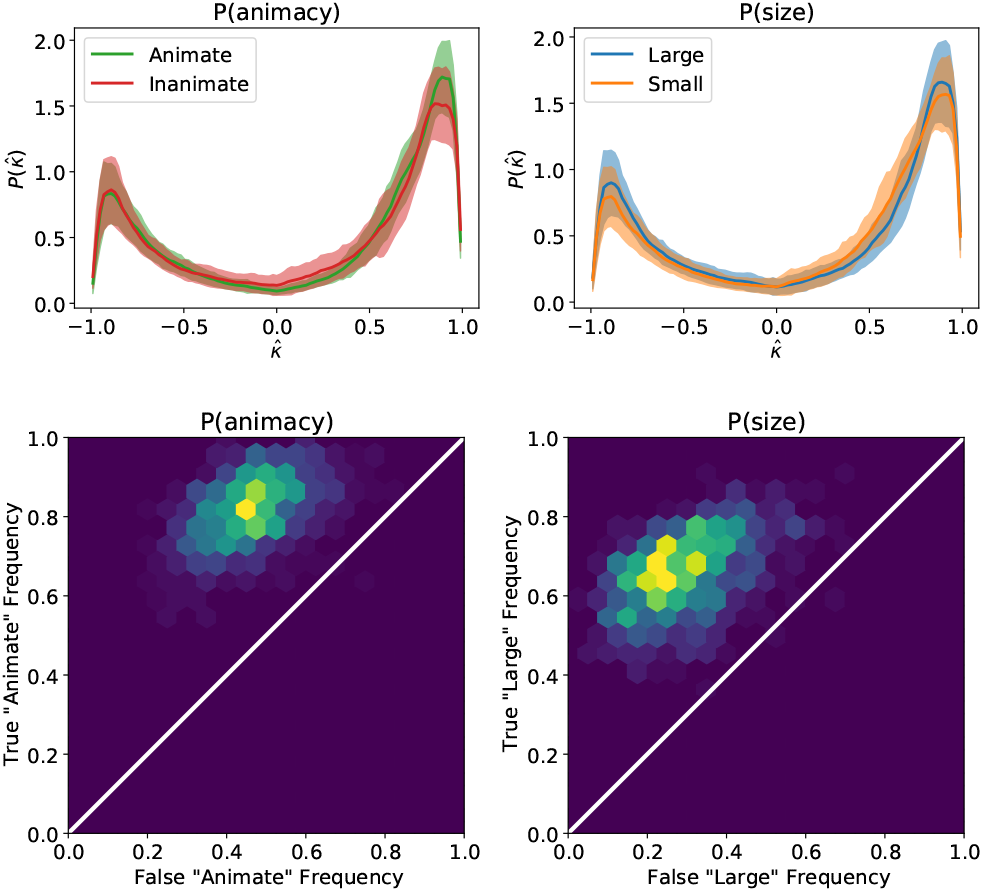
Similarly to Figure 3, the top two subfigures show the mean ±standard deviation of the NCC distributions for texforms corresponding to images of animate/inanimate and large/small objects (from [46]). The bottom two subfigures show the corresponding histograms of true positives and false positives from 1000 runs of a binary classifier (Eq. 3).

Overall, we see that the accuracy of the classifier for texforms is very similar to those of natural images; in fact, two of the categories yield marginally higher results. Additionally, Figure 8 shows false positive and false negative samples across each of the four categories. When classifying samples by animacy, we find that some images with considerable straight regions – such as the bull and bird – are confused for inanimate objects, while some highly textured samples – such as the stroller and wreath – are confused for animals. On the other hand, the samples mistakenly classified as small – such as the seal and armchair – are very rounded, contributing intermediate, positive curvatures, while the samples mistakenly classified as large - such as the Chihuahua and mouse running wheel – have a disproportionate amount of high or near-zero curvatures for their category.

**FIG. 8.**
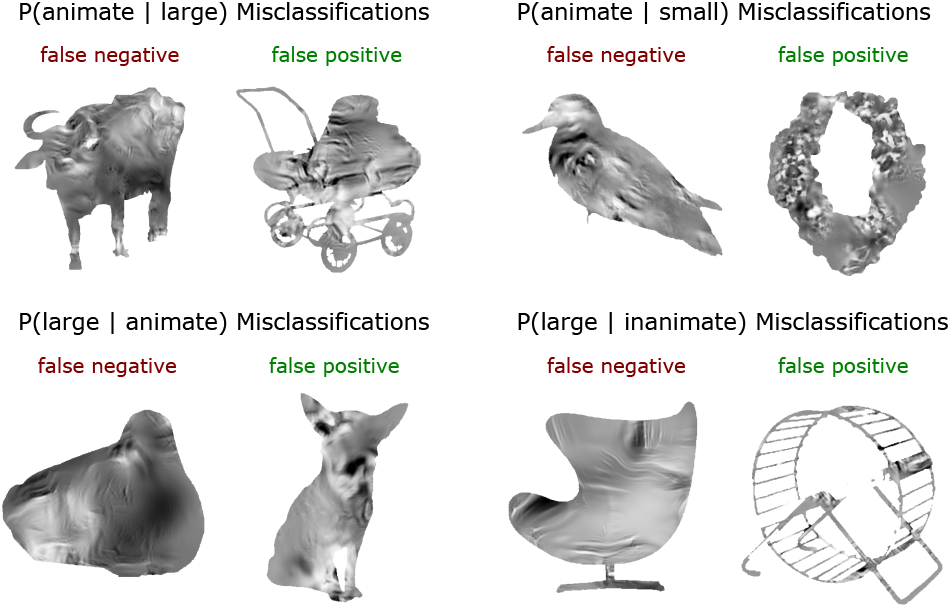
Following our analysis of natural images, we examine the errors made by a Bayesian classifier over the texform dataset (from [46]). We find similar qualitative characteristics in misclassified samples to those described in Figure 6: round large objects may be misclassied as small; inanimate objects with textured features may be misclassified as animate; animals are more difficult to classify by size.

## NCC DISTRIBUTION AS AN IMAGE FEATURE

Our results so far suggest that NCC statistics are consistent with experimental observations of cognitive categories across size and animacy, and thus raise the question of its use as an image feature in other downstream neural classification tasks. Complementing fMRI studies with human subjects, studies with macaque monkeys that directly measure neural activity in the brain have shown spatial localization and distinct topographical ordering of the neural responses in the visual cortex associated with the recognition of data types such as alpha-numeric characters in the Helvetica font, Tetris-like shapes, and simple cartoon faces [47], each of which are clearly distinguished by different geometric statistics, as shown in Fig. 9a,b along with their topographical ordering in the IT cortex. This raises the question: can normalized contour curvature reveal any structure across these categories? In line with standard image classification approaches, we can think of the calculated NCC distribution as a “feature vector” representing an image. Thus the NCC distribution can be combined with any other image features (such as average intensity, color information, etc.) by concatenation into a larger feature vector.

**FIG. 9.**
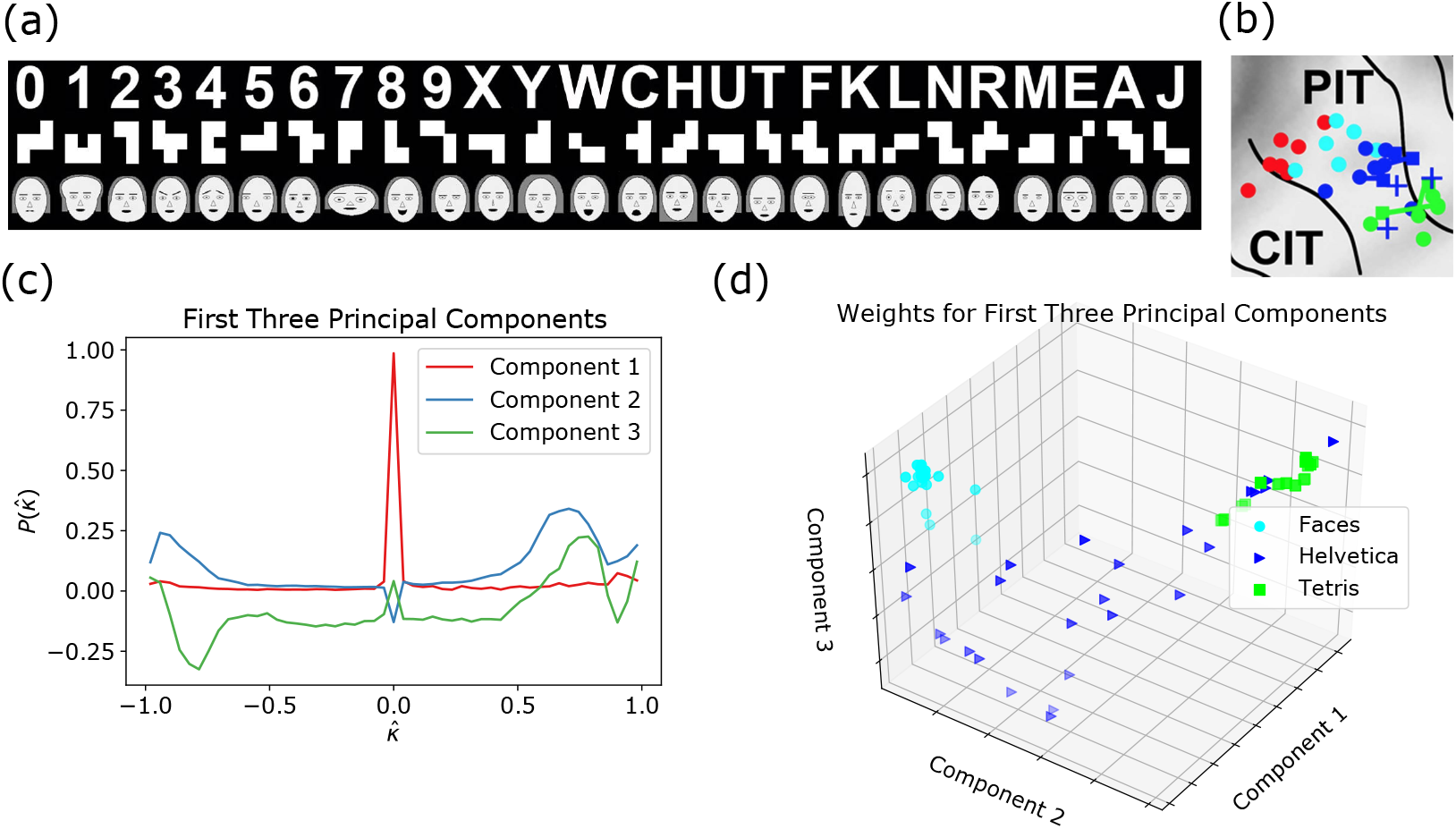
**(a)** Images of Tetris, Helvetica and cartoon face stimuli used by Ref. [47] to demonstrate proto-structure of the macaque IT cortex. **(b)** Spatial structure of neural activation measured in macaques in response to the stimuli, with cartoon faces in cyan, Helvetica in blue, tetris in green, and monkey faces in red (by Ref. [47]). **(c)** We apply principal component analysis of the NCC distributions across all stimuli. The first three components yield interpretation: Component 1 represents the amount of zero-curvature content, or straightness, Component 2 represents relative concave content, and Component 3 represents relative convex content. **(d)** By visualizing the 3D space of PCA coefficients corresponding to the first three components, we see clustering across categories consistent with the neural activation shown in (b).

### Methods

To visualize the relative clustering of NCC distributions across the different image categories, we project the features to a lower-dimensional space by applying principal component analysis (PCA). To carry out this approach, we construct a data matrix in which each column corresponds to an NCC distribution for a specific image, and calculate its singular value decomposition (SVD). Then, any NCC distribution can be approximated as a linear combination of orthonormal basis vectors (components). As we will see, the first three principal components have a natural geometric interpretation, and the projections of the data vectors into the three-dimensional space corresponding to these components allows us to evaluate separability across categories using the NCC as a feature vector.

### Results

In calculating the NCC for these, we use a relative filter size of *ρ* = 0.018 (as for the analysis of other images, the choice of this parameter does not change our results qualitatively). In Fig. 9c, we see clear structure in the principal components. In particular, the first component, which captures 89% of the variation in the data, represents a simple peak at a curvature of 0, therefore describing the amount of “straightness” in the data. The second component can be interpreted to represent concave content – from regions of the intensity surface which contain both positively- and negatively-curved contours – and the third component can be interpreted to represent convex content – which contains positive but not negative curvature content. This observation agrees with the finding that convexity and concavity drive differential neural responses to visual stimuli [48].

Visualizing the projections of the images onto the first 3 principal components shown in Fig. 9d, we see that the Tetris pieces with straight sides and corners, cluster tightly due to a high contribution from the first component. Similarly, the cartoon faces also cluster tightly due to their distinct structure – large positive curvatures (due to the round face), with small amounts of negative and zero-curvature content due to the eyes, nose, and other facial features. Finally, the Helvetica glyphs span the space between the latter two categories, as they contain a varied combination of straight rectangular portions and curved segments. In particular, a few are structurally very similar to “Tetris pieces” (such as the letters “I”, “L” and “H”). This ordering is particularly evident when we project the images onto the first two singular vectors; we see that it roughly matches the topographical organization observed in the brain (see Ref. [47], Fig. 3). Overall, we see that using the NCC statistics as a feature vector allows for linear separability of image classes in terms of a few geometrically interpretable principal components. **** We note that NCC permits interpretable geometric explanation for qualitative dimensions of “animate-inanimate” and “stubby-spiky” discovered by an artificial neural network in [49]: animate-inanimate distinctions are defined by zero-curvature content, while stubby-spiky distinctions are defined through high-curvature content.

## NCC AS A CLASSIFIER UNDER VARYING ILLUMINATION AND VIEWPOINT

Robust cognitive classification based on vision not only needs to be invariant to transformations of scaling, translation, and rotation, but also to variations in illumination and viewpoint. In the current context, we can ask if NCC-statistics based classifiers are robust to these variations? While it is difficult to provide theoretical guarantees in this context, we performed an experimental evaluation by quantifying the variation in NCC across illumination/viewpoint changes for individual objects, and comparing against inter-object variation using the Amsterdam Library of Object Images (ALOI) [50]. The ALOI is a dataset of 1000 household items, systematically photographed to vary viewing angle, lighting direction, and lighting color, for a total of 24 varied lighting conditions and 72 varied viewpoints, samples from which are shown on the left side of Fig. 10.

**FIG. 10.**
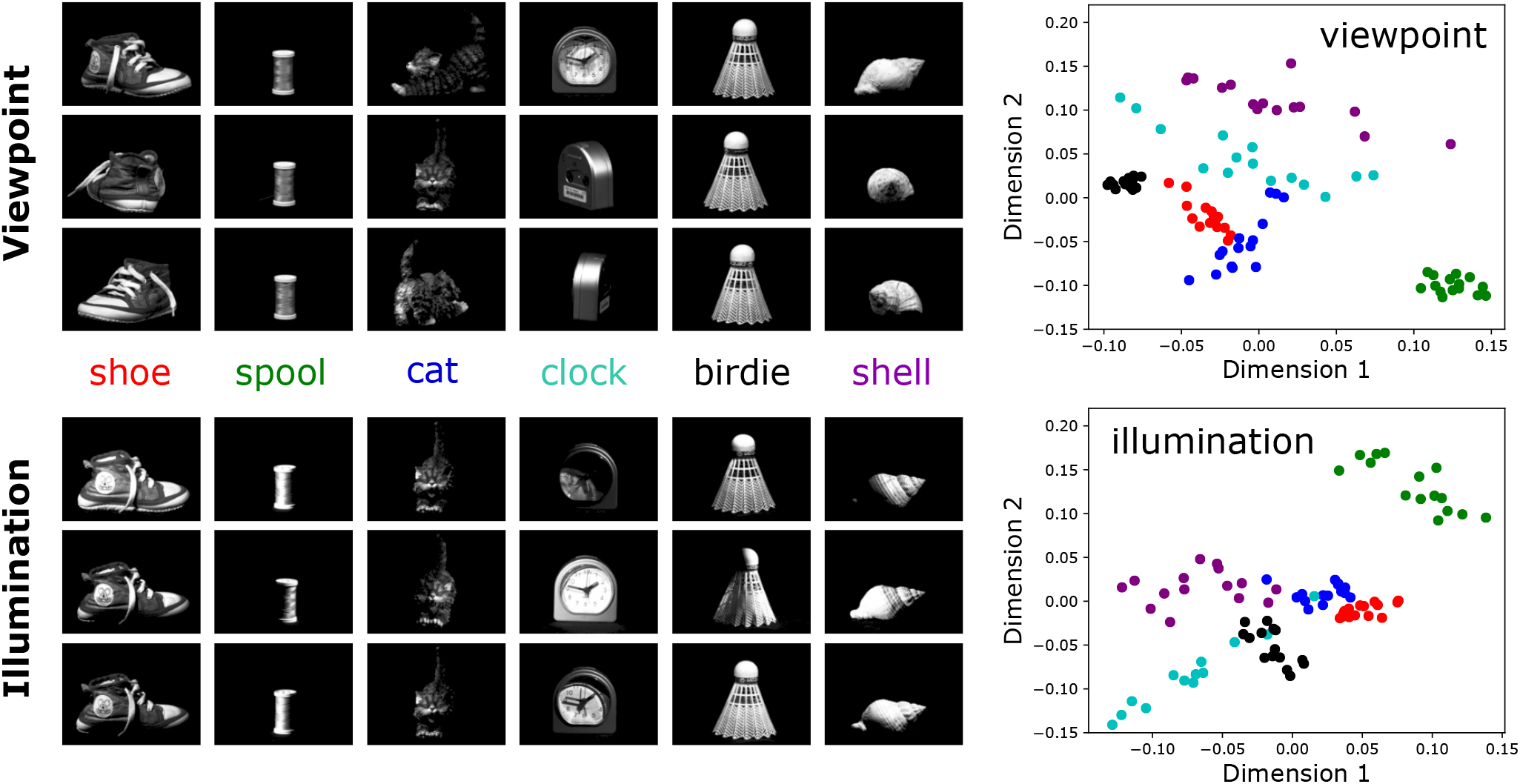
We examine robustness of NCC to two additional perceptual factors – viewpoint and illumination – by analyzing images of household objects taken under varying conditions from the Amsterdam Library of Object Images (ALOI) [50]. We consider six objects: a shoe, spool of thread, cat statue, small clock, tennis shuttlecock/birdie, and a seashell. After calculating NCC for images of the objects taken at varying conditions (varying viewpoint in the top row, or illumination in the bottom row), we project the NCC distributions to two dimensions through multi-dimensional scaling (MDS). The relative clustering of points in two dimensions can be considered a representation of the similarity of the NCC distributions for a given object. For instance, in the analysis of viewpoint variation, we find the birdie (black) and the spool of thread (green) have a very tightly clustered distribution, while objects such as the clock (cyan) and the seashell (purple) do not. This is understandable, as the former pair are radially symmetric, while the latter pair are difficult to recognize from the back. In the analysis of illumination variation, we find that reflective objects (such as the clock and shell) once again have more variance in the distributions, while objects with a more matte texture (such as the shoe and cat) have more consistent NCC.

### Methods

For our analysis, we choose six example object categories from the ALOI dataset, and randomly sample 15 images of varying lighting and viewpoint per object. To measure the similarity between probability distributions, we calculate the Jenson-Shannon divergence – a symmetric and smooth extension of the KL divergence – between NCC values for each pair of images. We then perform a multi-dimensional scaling (MDS) analysis to find a two-dimensional mapping which best preserves the “distances” between every pair of image samples, allowing us to visually evaluate inter- and intra-object similarity for varying illumination and viewpoint conditions.

### Results

In the right panels in Fig. 10, we show that the two-dimensional representations derived from NCC distributions represent consistent, and almost separable, clustering of images within object categories for variations in viewpoint and illumination. This is notable, as NCC is a simple metric which retains no correlative spatial information for an image. For example, in the case of illumination, there is larger variability for the seashell – for which some illumination angles cause part of the shell to fall into deep shadow, making it difficult to recognize – and the clock – in which the light casts shadows through the glass surface. In the case of changing viewpoint, we see very low variability within the spool of string and the shuttlecock, as these are radially symmetric, and higher variability in objects that look very different from behind, such as the cat and clock. All together, our results suggest that NCC is relatively robust to changes in illumination and viewpoint. Additionally, the spread of the clusters reflect sources of error that may be similar to those of human observers.

## A GENERATIVE MODEL FOR NCC DISTRIBUTIONS

Our use of the normalized contour curvature distribution as an cognitive classifier has highlighted how its invariance properties allow for an interpretable differentiation between image categories. We now turn to ask if it possible to create a generative model for constructing artificial images with specified NCC statistics related to known cognitive categories. To do so, we consider the theoretical distributions for the curvature statistics of a spatially varying Gaussian field with a correlation length distribution corresponding to a given NCC probability density, determined through an optimization procedure. This allows us to generate artificial images patch-by-patch with sampled correlation lengths, and thus compare the empirical, theoretical, and generated NCC statistics (see SI - Section F for more details on Gaussian random fields).

### Methods

We start with the assumption that Gaussian-filtered natural images can be roughly described in terms of distinct local patches drawn from Gaussian-correlated Gaussian fields with different correlation lengths *ξ*. Formally, this can be expressed as

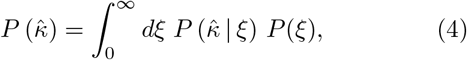

where 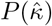 is the NCC distribution, *P*(*ξ*) is the correlation length distribution for a given cognitive category (e.g. animate) and 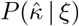 represents the single-pixel distribution for the NCC given that the local patch is described by a Gaussian-correlated Gaussian field with correlation length *ξ*. In discrete form, the probability distribution 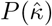 becomes the probability vector 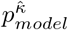, *P*(*ξ*) becomes *p*^*ξ*^, and 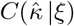 becomes Ξ, a matrix whose columns represent the NCC distribution for a Gaussian field with the corresponding correlation length *ξ*, such that

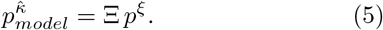

Formally, the elements of Ξ can be written in terms of the conditional cumulative distribution function 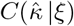 (See SI - Section G):

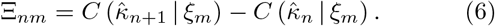

Then, we fit the correlation length probability vector *p*^*ξ*^ by taking it to be the probability vector minimizing the Kullback-Leibler divergence of the model distribution 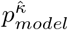 from the measured distribution 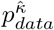,

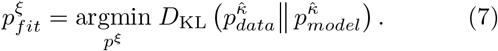

Note that this is a convex problem, which can be computed with standard optimization solvers (we used the “cvxpy” toolbox). Once we extract 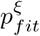, we can start generating artificial images. We first divide the image into patches of a chosen patch size. Then, for each patch, we draw a correlation length *ξ* from 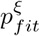, and fill in pixel values from a Gaussian-correlated Gaussian field with this correlation length, conditioning on pixels in the neighboring frames to maintain continuity across patches. Finally, once all panels have been filled in, we threshold the image, ignoring all pixels with intensities less than – *σ_f_* (where *σ_f_* is the standard deviation of the image).

### Results

In Fig. 11a, we show the fit for the correlation length distribution and see that it has a sparse three-peak structure. The first peak, occurring close to *ξ* = 0 is a boundary artifact, as calculated NCC distributions will have nonzero densities in the very first and last bin, which is theoretically impossible with nonzero correlation length. The second peak captures the majority of the distribution, while the last peak contributes to the bump at intermediate positive curvature values (around [0.5, 0.75]) – which, as we have argued earlier, is indicative of the circular characteristics of small objects. The relative weights associated with the peaks in the correlation length distribution allow *P*(*ξ*) to effectively capture the differences across both dimensions of animacy and size. Inanimate images have an additional peak at the maximum normalized correlation length, corresponding to a peak at 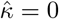, while large images have higher second peaks relative to third peaks, indicating greater probability density at the edges of the NCC distribution (see SI - Section E Fig. S7 for additional intuition and analysis of the structure of *P*(*ξ*)). In Fig. 11b we show that there is a good match between the NCC distribution of images from a category (e.g. animate), the theoretical distribution computed using the above optimization procedure, and the NCC of the images created using the generative algorithm. However, when the intensity fields associated with the generative approach are visualized, as shown in Fig. 11c, we note that they do not correspond to either natural images or even texforms. This is because the simple generative procedure captures only local – not global – spatial correlations, leading to generated images that exhibit smooth characteristics with irregular global geometric and topological structure, but do not represent individual objects.

**FIG. 11.**
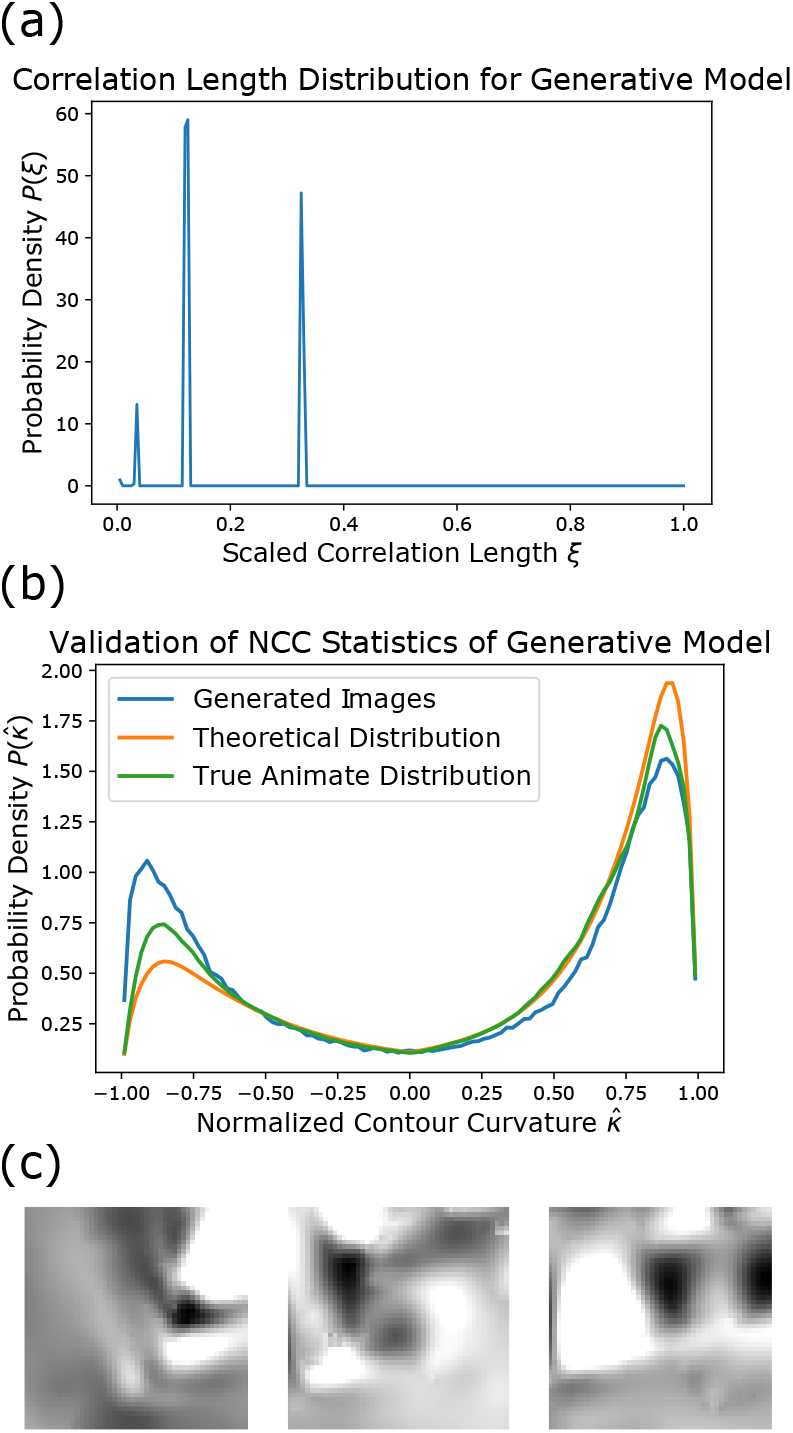
**(a)** Correlation length probability distribution for the animate image class extracted according to Eq. 7. **(b)** Comparison of animate NCC distribution, the model fit 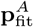 from Eq. 5 and the NCC distribution calculated from 100 thresholded (at an intensity of –*σ_f_*) artificial images. **(c)** Three images generated by constructing a Gaussian-correlated Gaussian random field with correlation length drawn from the distribution in (a).

## DISCUSSION

While many modern neural network based approaches are remarkable in their ability to categorize visual stimuli, there remains a need for simple, interpretable, properly invariant metrics to describe cognitive categorization, because the latter is more likely to illuminate the underlying mechanisms in both artificial neural networks and the neural processing behind human cognition. Our use of the normalized contour curvature distribution aims to rejuvenate a geometric idea in a probabilistic setting, recognizing that we need metrics that are invariant to the Euclidean group of rotations and translations, and intensity scaling, but also accounting for statistical variability within and across images. We have shown that simple metrics based on the NCC distribution are consistent with multiple experimental findings in humans and monkeys; it can distinguish between cognitive categories, is robust to changes in lighting conditions and viewpoint, and has plausible implementability in neural circuitry given that curvature has roots in characterizing orientational information, and can then be pooled.

One limitation of the proposed NCC distribution as a metric is that it does not account for the spatial distributions of curvature content. When perceiving complex natural scenes, or even more detailed objects, we clearly rely on non-local and/or higher-order statistics of the visual field. To mitigate this, future development of this metric could integrate NCC with a pyramid framework, in which a feature descriptor is applied at varying locations and scales [51]. However, it has also been shown that the neural response to an image is not distinct. By estimating the size of receptive fields it is possible to construct artificial images with distorted peripheral intensities which are perceptually indistinguishable for a human observer - metamers [52], which then led to alternative artificial stimuli, such as texforms [46, 53] which we also considered here. This coarse-graining of spatial information suggests that local pooling of neural activation is an important aspect of image processing, consistent with our results that show that the NCC distributions of modified texforms contain similar defining curvature characteristics across animacy and size as in natural images. Finally, our generative model for NCC highlights a particularly difficult problem by failing to capture global geometric and topological considerations in images; how we might augment a statistical-geometric approach such as the one used here with ideas from integral geometry [54] remains an open question.

## ACKNOWLEDGMENTS

We thank Marge Livingstone for asking a question that launched this study, Talia Konkle for providing us with the animate/inanimate and large/small image sets and both of them for discussions, the Harvard Quantitative Biology Initiative and the NSF-Simons Center for Mathematical and Statistical Analysis of Biology at Harvard, award no. 1764269 (I.T., L.M.), and the Henri Seydoux Fund for partial financial support. All code and data for these calculations is available at the GitHub repository XXX

## Notes

### Competing Interest Statement

The authors have declared no competing interest.

